# DNA methylation regulates a key sex differentiation gene in *Pogona vitticeps*, a dragon lizard with sex reversal

**DOI:** 10.64898/2026.06.15.731764

**Authors:** Benjamin J. Hanrahan, Susan Wagner, Nicholas C. Lister, Sarah L. Whiteley, Lei Xiong, Arthur Georges, Paul D. Waters

**Author notes:** Correspondence: Paul D. Waters, Arthur Georges.

## Abstract

During embryonic development bipotential gonads differentiate into either testes or ovaries under the direction of mutually exclusive gene networks. DNA methylation has been shown to regulate gene expression and to play a role in many developmental processes. This includes sex differentiation in which DNA methylation is linked to control of key sex genes. Whether DNA methylation regulates sexual differentiation in species where environmental influences such as temperature are involved is less clear. We conducted a genome-wide study in embryonic gonads of the central bearded dragon (*Pogona vitticeps*), a lizard with temperature induced sex reversal, and compared DNA methylation patterns to gene expression profiles at a stage of early sex differentiation. Overall, sex reversed ZZf females were found to have lower global methylation than both canonical sexes, ZZm males and ZWf females. We found that the expression of a key gene in sex differentiation, *Amh*, is regulated via DNA methylation. *Amh* is a driver of testis differentiation and is repressed in genetically determined females, as well as in temperature sex-reversed females, by hypermethylation around its transcription start site. Although the trigger of ovary determination is different in both groups of females, one by genetic complement and one by a temperature signal, downstream regulation of gene expression converges and seems to follow similar mechanisms.

## INTRODUCTION

Sexual differentiation results in two discrete outcomes, development of a testis or ovary, directed by mutually reinforcing gene networks that also suppress the opposite pathway (Nagahama et al. 2021). DNA methylation, the addition of a methyl group to cytosine residues, provides a layer of gene regulation, often repressing transcription when present in promoters and enhancers (Head 2014; Schübeler 2015).

Increasing evidence shows that DNA methylation fine-tunes sex differentiation pathways and contributes to cross-program suppression in vertebrates with both genetic and environmental sex determination (ESD) (Navarro-Martín et al. 2011; Piferrer 2013; Liu et al. 2022). Examples from mammals, birds, fish, and reptiles demonstrate that methylation of key sex-related genes (e.g. *Sry*, *Sox9*, *Cyp19a1*, *Amh*) influences whether gonads develop as testes or ovaries (Ellis et al. 2012; Okashita et al. 2019; Dong et al. 2020).

In species with ESD, DNA methylation is becoming a particularly important avenue of research into how environmental cues such as temperature can influence sexual fate (Navarro-Martín et al. 2011; Matsumoto et al. 2013; Lin et al. 2021; Liu et al. 2022). Key sex-related genes such as *Sox9*, *Cyp19a1* (also known as *aromatase*) and *Amh* have shown differential methylation associated with changes in gene expression in reptile species with ESD (Parrott et al. 2014; Matsumoto et al. 2016; Dong et al. 2020). Particularly in the gonads or during early embryonic development, these patterns suggest a role for DNA methylation in locking-in environmental signals to influence sex differentiation. As such, in reptiles where temperature can influence sexual outcomes, methylation is a candidate mechanism for translating thermal cues into gene expression changes (Parrott et al. 2014; Matsumoto et al. 2016).

The central bearded dragon (*Pogona vitticeps*) combines a ZZ/ZW genetic system with temperature-induced sex reversal, producing ZZ females at high incubation temperatures (Ezaz et al. 2005; Quinn et al. 2007; Holleley et al. 2015). The interaction of temperature and genetic sex determination in *P. vitticeps* presents an opportunity to study the effect environment can have on sex via comparison of sex reversed and non-sex reversed individuals. Previous work on both understanding the genetic regulation of sex determination in the bearded dragon (Whiteley et al. 2021; Wagner et al. 2023) and the mechanisms by which temperature-induced sex reversal occurs (Deveson et al. 2017; Whiteley et al. 2022) have suggested an intricate system of genetic control for which much remains unknown. Recent work has also identified duplications of *Amh* and *Amhr2* on the Z chromosome and isoform differences in *Nr5a1* between ZZ and ZW individuals, which highlights the complex genetic regulation of sex in this species (Zhang et al. 2022; Guo et al. 2025; Patel et al. 2025).

Here, we present the first whole-genome methylation analysis for *Pogona vitticeps,* a reptile with temperature-induced sex reversal. By combining methylomes and transcriptomes of stage-12 embryonic gonads, we reveal the epigenetic regulation of key sex differentiation genes, focusing on *Amh* and its Z-linked copy *Amh-Z*, as well as *Amhr2* and *Amhr2-Z*. This provides new insight into the molecular regulation of temperature-induced sex reversal and has a broader impact on our understanding of temperature on reptile sex determination.

## RESULTS

### Females are hypomethylated compared to males

To understand the role of epigenetics in sex differentiation of *P. vitticeps*, we employed whole genome bisulphite sequencing and RNA-sequencing (Whiteley et al. 2021) to correlate DNA methylation and gene expression in developing gonads. We used embryonic stage 12 gonads, when they are in an early phase of differentiation (Whiteley et al. 2017; Whiteley et al. 2018), and compared concordant males (ZZm), concordant females (ZWf) and temperature driven sex-reversed females (ZZf).

Clustering analysis and principal components analysis (PCA) showed that the methylomes of samples clustered with their sex condition (ZWf, ZZf, and ZZm) (**Figure S3A, B**). The ZZf samples, which were incubated at 36°C, displayed a larger distance to the other two clusters, which shared a common incubation temperature of 28°C (**Figure S3**).

ZWf females were significantly hypomethylated compared to males (**Figure 1A**), with global hypomethylation most pronounced in gonads from sex-reversed ZZf females (median methylation rate: ZZm 73.0 %, ZWf 71.4 %, ZZf 68.0%). Between all three groups, 4-8% of 1 kb windows were significantly differentially methylated (q-value < 0.001; **Figure S3C-E**). A higher number of differentially methylated regions (DMRs) were hypomethylated in both ZZf and ZWf females compared to ZZm males (**Figure 1B, D**). This was prominent in sex-reversed ZZf females, which had six times more hypomethylated DMRs than hypermethylated DMRs compared to both ZWf and ZZm (**Figure 1C, D**).

**Figure 1.**
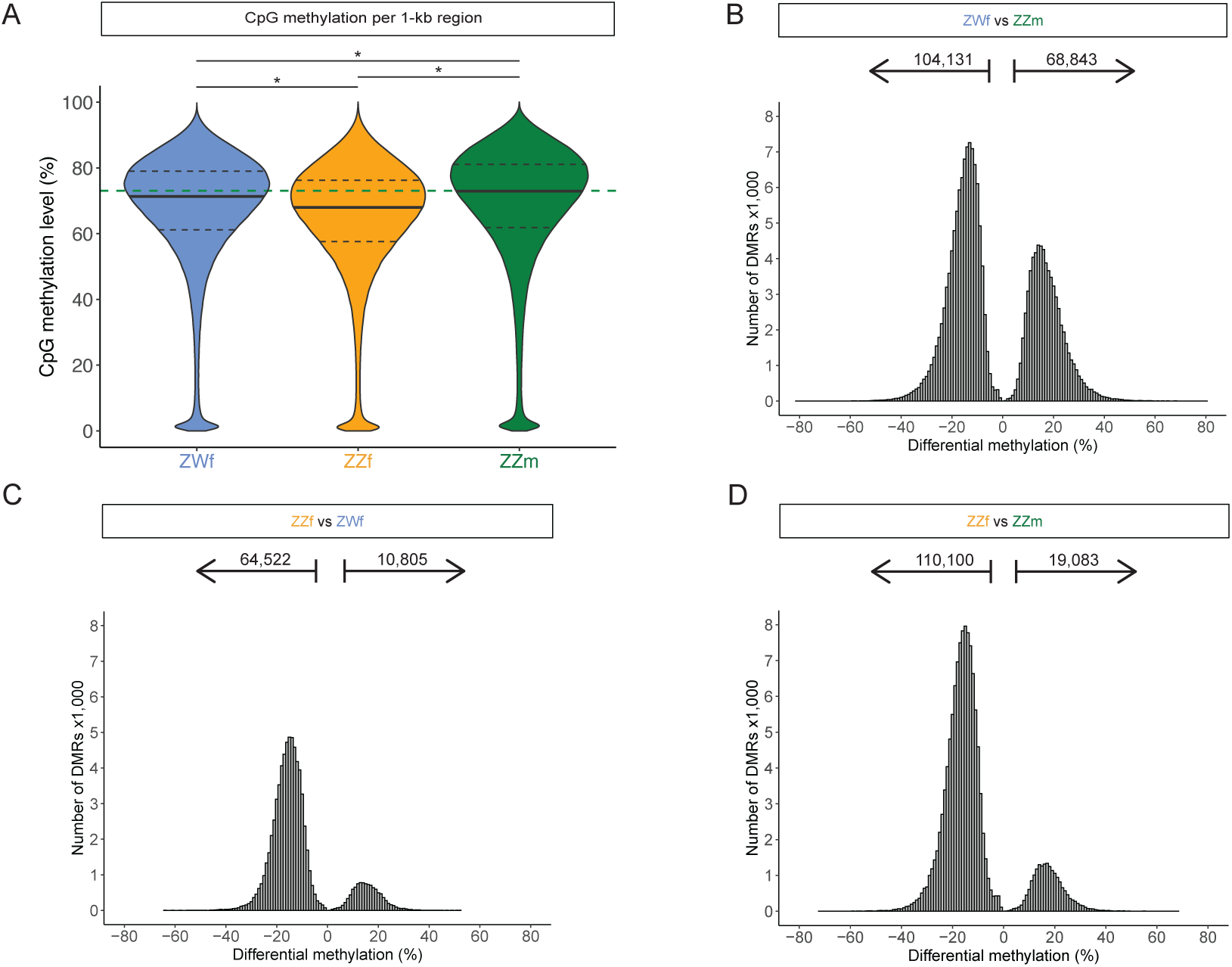
Hypomethylation in canonical females and sex reversed females. (A) Median CpG methylation levels in 1 kb windows for each group (ZWf, ZZf, ZZm). Medians are shown as a solid black line and quartiles as dotted black lines. Only windows with a CpG read depth of at least 20 in all samples were included (770,741 regions). Green dashed line is the ZZm median to facilitate comparison across groups. * indicate a p-value < 0.0001 (Kruskal-Wallis and Conover-Iman tests). (B-D) The distribution of significantly differentially methylated regions (DMRs, q-value < 0.01) is plotted as a histogram for three pairwise comparisons: (B) ZWf v ZZm, (C) ZZf v ZWf, (D) ZZf v ZZm. The numbers represent how many DMRs have significantly increased and decreased methylation levels.

### DNA methylation levels negatively correlate with transcription

To investigate the impact of DNA methylation on gene expression, we assessed the DNA methylation levels across the transcription start sites (TSSs) of genes. Decreased methylation at TSSs correlated well with high or very high gene expression levels. The median methylation levels of lowly expressed genes were approximately 10% higher than highly and very highly expressed genes directly over the TSS. Therefore, a relatively small change in methylation levels correlated with a large change of gene expression. Genes that had very low or no expression at all displayed high methylation levels over the entire TSS region (**Figure 2A**).

**Figure 2.**
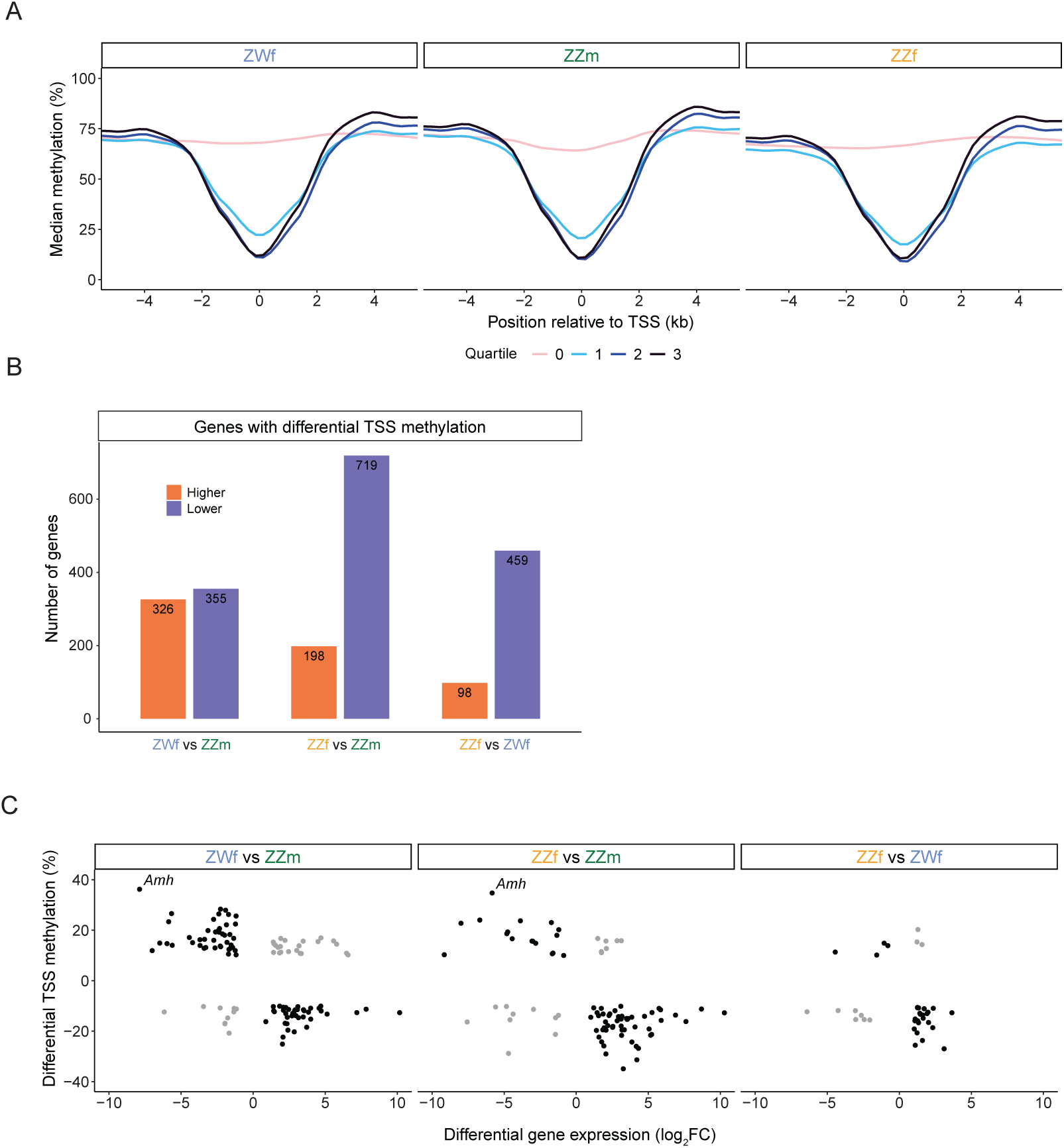
Metagene analysis of DNA methylation levels across all TSSs and the TSS of *Amh*. (A) CpG methylation levels across the transcription start sites (TSSs) of all genes for each sex: ZWf, ZZf and ZZm. Genes were divided into expression quartiles: very high, high, low, very low/no expression. The smoothed lines represent median methylation at each position relative to the TSS ±5 kb with line colour indicating expression quartiles. (B) Number of genes with differential TSS methylation for each pairwise comparison (±500bp of the TSS). Column colour indicates whether a TSS had increased or decreased methylation. (C) For DEGs that also have a DMT the differential methylation rate as a percentage is plotted against the differential gene expression (log_2_FC). The expected negative correlations between methylation and expression are indicated by black points, whereas the unexpected positive correlations are indicated by grey points.

CpG methylation levels approximately 4 kb upstream of genes in all expression bins were similar. In contrast, regions 4 kb downstream of high and very highly expressed genes displayed higher methylation levels (**Figure 2A**). These patterns were consistent across the three genotypic and phenotypic groups (ZWf, ZZf and ZZm).

### Upregulated gene expression in sex-reversed females correlates with TSS hypomethylation

Because metagene analysis revealed a negative correlation between the DNA methylation level at TSSs and expression levels of associated genes, we questioned if regions ±500 bp of TSS displayed differential methylation. Both ZZ and ZW females had more than 650 genes with a differentially methylated TSS (DMT) when compared to males (ZZf v ZZm: 681 DMTs, ZWf v ZZm: 917 DMTs). Between sex-reversed ZZf and canonical ZWf there were 557 DMTs (**Figure 2B**), with a split of three times more DMTs being hypomethylated than hypermethylated (**Figure 2B**).

Between 5 - 12% of the genes with a DMT were also significantly differentially expressed (adjusted p-value < 0.05, fold change > 1.5) (**Figure 2C**). Most genes (182 of 241) displayed the expected negative correlation of gene expression level and DNA methylation level (**Figure 2C**). Gene ontology (GO) analysis of the genes with the expected negative correlation between TSS methylation and expression in the ZZf v ZZm comparison, showed “metal ion transport” and “monoatomic cation transport” as enriched biological processes, as well as terms related to regulation of development (**Table S1**).

Genes identified in the GO analysis of particular interest included genes related to sexual development and regulation; *Amh* and *Gata1,* as well as genes involved calcium transport and signalling; *Cacnb3* and *S100a1* (**Table S1**). Other, genes of interest identified to be differentially expressed with a DMT in the ZZf v ZZm comparison include *aromatase* (*Cyp19a1*, here identified as *LOC110077216*), which is involved in estrogen production. A further two genes encoding estrogen receptors, *Esr1* and *Greb1* were also differentially expressed with a DMT (**Table S2**).

### *Amh* genes and receptors

*Amh* is a gene that plays a major role in sex differentiation. It was differentially expressed between all comparisons and had a differentially methylated TSS. Both sex-reversed ZZf females and concordant ZWf females had higher DNA methylation levels around the TSS of *Amh*, which correlated with lower expression compared to ZZm males (**Figure 3 A, B**). A paralog of *Amh* (called *Amh-Z*) was recently identified on the Z chromosome (Guo et al. 2025; Patel et al. 2025). Methylation levels over the *Amh-Z* TSS showed no difference between sexes, there was a drop in methylation at the TSS in all sexes characteristic of active gene expression (**Figure 3C)**. Despite the same methylation profiles in all sexes, there was higher expression in ZZm compared to both ZZf and ZWf females (**Figure 3D**).

**Figure 3.**
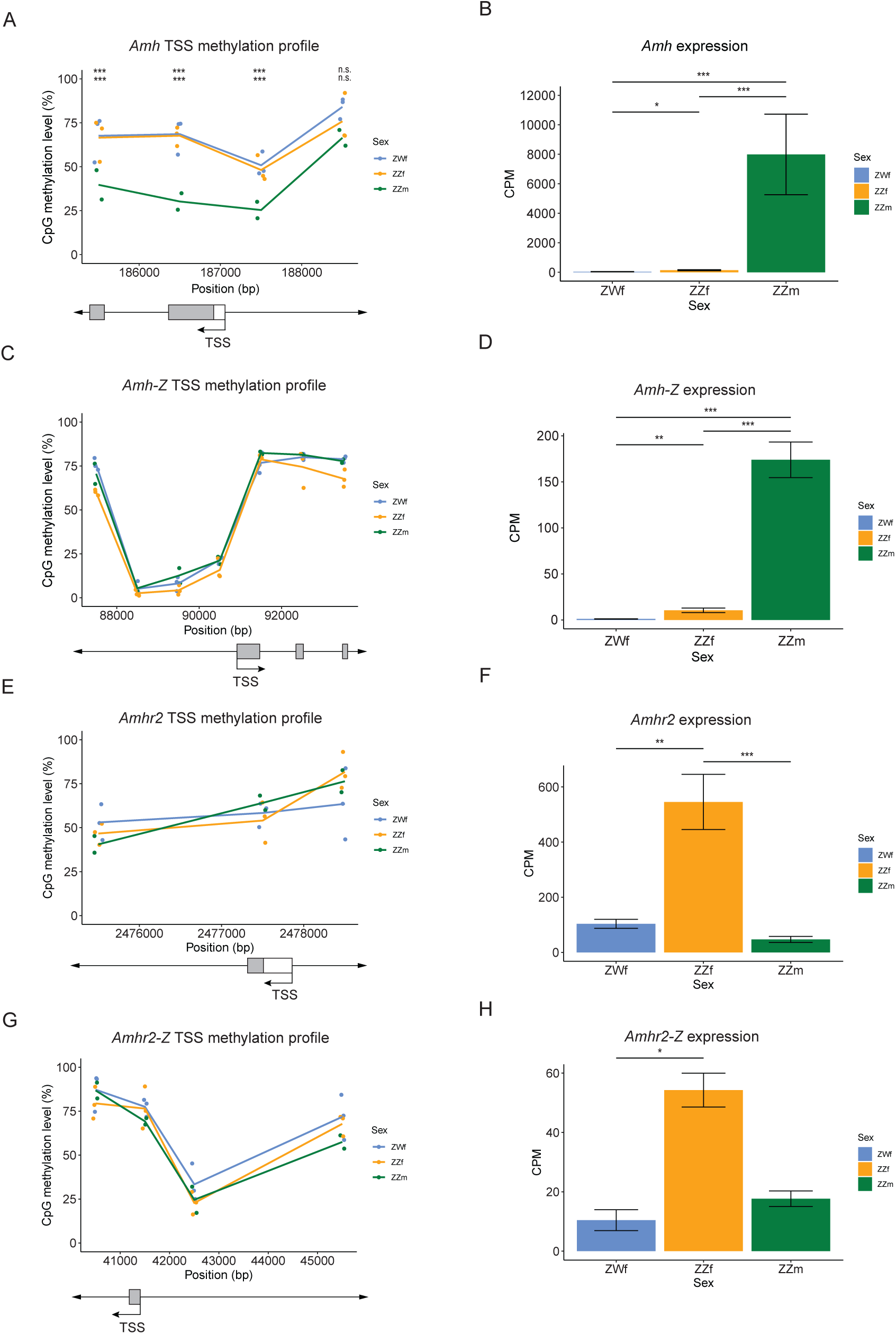
TSS DNA methylation profiles and expression levels for *Amh* genes and its receptors. DNA methylation profile around TSSs (left) and gene expression (right) for the genes: *Amh* (A, B), *Amh-Z* (C, D), *Amhr2* (E, F) and *Amhr2-Z* (G, H). (A, C, E, G) DNA methylation levels in 1 kb windows were plotted either side of the transcription start sites. Points for each sample were plotted with a line joining the means of each sex. Sex is indicated by point and line colour. (B, D, F, H) Gene expression is plotted in counts per million (CPM) for each sex. Mean expression is shown as bar plots; error bars represent standard error. ***, **, * indicates significantly different expression between sexes (*** = FDR < 0.001,** = FDR < 0.01 * = FDR < 0.05).

We also investigated the DNA methylation profile and gene expression level of the gene encoding the receptor for AMH, *Amhr2* (**Figure 3E, F**). No difference in methylation was identified over this TSS between the sexes. However, ZZf females show significantly higher expression compared to both ZZm males and ZWf females (**Figure 3F)**. *Amhr2* also has a paralog on the Z chromosome, *Amhr2-Z*, which showed a similar expression pattern to *Amhr2.* The DNA methylation profile over the TSS of *Amhr2-Z* did not show a difference between sexes (**Figure 3G**). However, ZZf females had higher gene expression compared to ZWf females, but the difference to ZZm males was not significant (**Figure 3H**).

## DISCUSSION

Here we examine the genome wide DNA methylation patterns in the developing gonads of the central bearded dragon, *P. vitticeps*. We found that concordant ZWf and sex-reversed ZZf females have lower overall methylation levels compared with ZZm males. ZZf females and ZWf females also had a higher number of DMRs with lower methylation when compared to ZZm males. We then focussed on methylation flanking the TSSs of genes. By grouping genes, with the expected negative correlation of methylation around the TSS and gene expression in matched gonad samples, we observed an enrichment of genes related to GO terms relevant to a current model of sex reversal (Castelli et al. 2020). We also, identified that the key sex determining gene *Amh* had lower expression in both ZWf and ZZf females in comparison to ZZm males, coupled with higher DNA methylation directly over the TSS in both female sexes. This suggests DNA methylation is repressing expression in this key male determining gene in both concordant and sex reversed female *P. vitticeps*.

Overall, concordant females (ZWf) of *P*. *vitticeps* had lower global methylation than concordant males (ZZm) (**Figure 1A**). This pattern of lower methylation in female gonads is observed in several fish species, including zebrafish (Laing et al. 2018), bluehead wrasse (Todd et al. 2019) and half-smooth tongue sole (Shao et al. 2014). In the case of sex reversed *P. vitticeps*, ZZf females had lower genome wide methylation (68 %) than both ZZm males (73 %) and ZWf females (71.4 %, **Figure 1A**).

In the tiger pufferfish, a fish species with sex reversal, global DNA methylation does not show a clear difference between sexes (Zhou et al. 2019). In another fish, the half-smooth tongue sole, where incubation at high temperatures causes the sex reversal of ZW individuals to ZWm males, there is a higher overall methylation level in the sex reversed ZWm males and ZZm males (78%) compared to ZWf females (67%) (Shao et al. 2014). The lower methylation observed in sex reversed ZZf females in *P. vitticeps* may be indicative of a genome-wide change resulting from high temperature incubations which cause sex reversal.

Gene ontology analysis of the genes with DMTs and differential expression with the expected negative relationship in the ZZf female to ZZm male comparison identified “metal ion transport” and “monoatomic cation transport” as significantly enriched biological processes. Included in these sets was the gene *Cacnb3*, a protein that is a subunit of a plasma membrane channel controlling the passage of Ca^2+^ (Collin et al. 1994; Arikkath and Campbell 2003). *Cacnb3* plays a role in mediating the homeostatic response to high amounts of Ca^2+^ in the cell, which results in regulation of Ca^2+^ signalling pathways (Namkung et al. 1998; Murakami et al. 2000).

The gene *S100a1* was also identified in the GO analysis. It had lower gene expression and higher TSS methylation in ZZf females compared to ZZm males. The function of the encoded protein involves binding free Ca^2+^ ions and facilitating Ca^2+^ movement (Kiewitz et al. 2003; Wright et al. 2005).

A model for how temperature causes sex reversal in *P. vitticeps* suggested that increases in cellular Ca^2+^ may be involved. This model was based on a hypothesis that cellular calcium and reactive oxygen species capture the temperature signal that causes sex reversal (Castelli et al. 2020). In stage 6 embryonic gonads, Ca^2+^ signalling related genes (including *Cacnb3*) are upregulated in ZZ females compared to ZW females (Whiteley et al. 2021). In our stage 12 gonads *Cacnb3* had higher expression and lower TSS methylation in ZZ females than in both canonical sexes (i.e. ZW females and ZZ males). This DNA methylation difference indicates that it may be important in regulating the expression of *Cacnb3* and *S100a1* in *P. vitticeps* sex reversal.

A screen of genes with differential methylation around their TSS and with differential expression identified the candidate sex determining gene *Amh,* which was also identified in several GO terms (**Table S1**). In the promoter and TSS there was low DNA methylation and relatively high expression in ZZm males. In contrast there was high DNA methylation and low expression in ZZf and ZWf females (**Figure 3 A,B**). We have previously shown that *P. vitticeps* has a unique expression profile of *Amh* compared to mammals (Wagner et al. 2023). This includes females showing some expression of *Amh* in developing gonads. In mammals there is no expression of *Amh* in female gonads (Rey et al. 2003). The methylation pattern observed here of high methylation in females (both genotypes) and lower DNA methylation in males provides a candidate epigenetic mechanism for which *Amh* is silenced in *P. vitticeps*. The high DNA methylation in sex reversed ZZf females, which had lower *Amh* expression than ZWf females, suggests that it is an important factor for female development.

We also examined the gene expression and DNA methylation profile of a recently identified copy of *Amh* in the non-recombining region of the Z chromosome (annotated in the Pvi1.1 genome as *LOC110070405*), referred to here as *Amh-Z* (Guo et al. 2025; Patel et al. 2025). The gene expression follows that of the autosomal *Amh* in the stage 12 embryonic gonads, with higher expression in ZZm males compared to both ZZf and ZWf females. The methylation profile of *Amh-Z* has reduced DNA methylation near the TSS for all sexes, which does not indicate that there is sex specific regulation by DNA methylation of this *Amh* copy.

Differential expression was also observed for the gene *Amhr2*. However, in contrast to *Amh*, sex reversed ZZ females had higher expression than both ZZ males and ZW females (**Figure 3F**). *Amhr2* encodes the sole receptor for the AMH hormone, which in mammals facilitates mullerian duct regression (Dyche 1979; Allard et al. 2000). In chicken, *Amhr2* also has an important role in testis formation (Cutting et al. 2014). As *Amhr2* expression is associated with early male development in both mammals and birds its higher expression in sex reversed ZZf females is surprising. Like *Amh*, a copy of *Amhr2* occurs in the non-recombining region of the Z chromosome (referred to here as *Amhr2-Z*), which also had higher expression in ZZf females than ZWf females (**Figure 3H**). This indicates that *Amhr2* may play a different role in gonadal development of *P. vitticeps* and the sex reversal of ZZf females, or that high expression levels of *Amhr2* in males occur at a later developmental stage.

Epigenetic mechanisms in the form of chromatin modifiers and their machinery have previously been implicated in sex determination and sex reversal in *P. vitticeps* (Deveson et al. 2017; Whiteley et al. 2022). Here, we provide an added layer of epigenetic complexity in DNA methylation and, for the first time, examine the methylome of a sex reversing reptile via whole genome bisulphite sequencing. We identify *Amh* as a key sex determining gene being regulated by DNA methylation, which may be the mechanism by which the *Amh* expression profile is locked in during embryonic gonad development.

## MATERIALS AND METHODS

### Experimental design

Our experimental setup used one gonad each for the DNA methylome and the RNA transcriptome, this allowed us to assess the correlation between the levels of DNA methylation and gene expression.

### Animal breeding, egg incubations and embryo sampling

Animal breeding, egg incubation and embryo sampling was performed as described in Whiteley et al. (2021). Here repeated, eggs were obtained during the 2017-18 breeding season from the research breeding colony at the University of Canberra. Breeding groups comprised three sex reversed females (ZZf) to three males (ZZm), and three concordant females (ZWf) to two males. Paternity was confirmed by SNP genotyping (Whiteley et al. 2021). Females were allowed to lay naturally, and eggs were collected at lay or within two hours of lay. Eggs were inspected for viability as indicated by presence of vasculature in the egg, and viable eggs were incubated at 28°C in temperature-controlled incubators (±1°C) on damp vermiculite (4 parts water to 5 parts vermiculate by weight). Clutches from sex reversed females (that is, ZZf x ZZm crosses) comprised eggs with only ZZ genotypes. These were initially incubated at 28°C to entrain and synchronise development. After 10 d of incubation, half of the eggs selected at random from each clutch was shifted to 36°C. Clutches from ZWf x ZZm crosses were incubated at 28°C throughout the incubation period. Eggs were sampled at times corresponding to developmental stage 12, equating a recently differentiated gonad (Whiteley et al. 2017; Whiteley et al. 2018). Gonads were sampled from 3 ZWf females, 2 ZZm males and 3 ZZf females. Refer to **Table S3** for a summary of information and data accessibility for all samples. Embryos were euthanized by intracranial injection of 0.1 ml sodium pentobarbitone (60 mg/ml in isotonic saline). Individual gonads were dissected from the mesonephros under a dissection microscope and snap frozen in liquid nitrogen. Embryos were genotyped using previously established protocols (Holleley et al. 2015; Whiteley et al. 2017). Briefly, this involved obtaining a blood sample from the vasculature on the inside of the eggshell on an FTA® Elute micro card (Whatman). DNA was extracted from the card following the manufacturer protocols, and PCR was used to amplify a W specific region (Holleley et al. 2015) so allowing the identification of ZW and ZZ samples.

### DNA extraction and whole genome bisulphite sequencing for methylome analysis

One gonad of the pair was used for DNA extraction and bisulfite sequencing. Bisulfite conversion and library preparation was performed using the Zymo Research Pico Methyl-Seq Library Prep Kit (Cat. No. D5456). The protocol was carried out as per manufacturers instruction with the exception of Section 4: Step 2. Here, a total of 10 amplification cycles were used (instead of 6) as recommended for low input samples. The libraries were sequenced on the Illumina NovaSeq 6000 platform at the Ramaciotti Centre for Genomics (UNSW Sydney, Australia) producing 100 bp paired-end reads with an average of 420 million read pairs per sample.

### Data analysis for DNA methylome profiles

Fastq files were assessed with FastQC (Andrews 2010) and trimmed accordingly with trimmomatic v0.38 (Bolger et al. 2014) using the following options: ILLUMINACLIP:TruSeq3-PE.fa:2:30:10:2:TRUE HEADCROP:15 SLIDINGWINDOW:4:15 MINLEN:36. Paired forward and reverse reads were mapped to indexed *P. vitticeps* genome assembly pvi1.1 (GCA_900067755.1) (Georges et al. 2015) with bismark v0.22.3 (Krueger and Andrews 2011) using the following options: --non_directional -D 20 -R 3 - N 0 -L 20 --score_min L,0,-0.6. Reads that aligned to the same position and strand in the genome were thought to be PCR duplicates and bismark function deduplicate_bismark. Genome-wide CpG methylation reports were extracted with the bismark function bismark_methylation_extractor using the following options: --scaffolds --cytosine_report. Cytosine reports were loaded into R v4.2.3 for further analysis. All analyses of methylation data were carried out with the R package methylkit v1.22.0 (Akalin et al. 2012) using custom scripts. Read depths for all CpGs within 1-kb regions of the genome were cumulated into windows. Scaffolds smaller than 1 kb were therefore excluded. For further analysis, regions with a cumulated CpG read depth of less than 20 in any of the eight samples were excluded, resulting in 770,741 windows. The calculateDiffMeth() function was used to determine which of the 1 kb windows were differentially methylated in a pairwise fashion between the three sexes. A q value < 0.01 defined a window as being differentially methylated.

### DNA methylation profiles around transcription start sites

TSSs were clustered into four groups according to the expression level of the genes, as described below in the paragraph ‘Data analysis for transcriptome profiles’. The cytosine reports generated from Bismark were imported into R using the methRead function with a minimum coverage of 1 and normalizeCoverage was used to normalise coverage between samples. The unite function was used to unite and destrand samples. Percentage methylation for each position was calculated using the percMethylation function.

The methylation dataframe was then intersected with start and end positions calculated as 5 kb either side of the TSSs of genes as annotated by the Pvi1.1 genome. Sites within the 10 kb window were retained and the relative position to the TSS was calculated. The methylation and expression group data frames were merged to determine methylation of genes belonging to each group. For each expression group the median of each site across the 10 kb window around the TSS of all genes in that group was calculated and plotted as a smoothed line (**Figure 2A**).

### Differentially methylated TSSs (DMTs)

Figure 2A indicated that the largest point of difference in TSS methylation between highly and lowly expressed genes was around a 1 kb window (±500bp) centred on the TSS. As such, to determine differential TSS methylation, cytosine reports were imported into R and normalised as above. Total methylation in a 1 kb window around each TSS as annotated in the Pvi1.1 genome was then calculated using the regionCounts function from methylKit in conjunction the package GenomicRanges v1.50.2 (Lawrence et al. 2013). The calculateDiffMeth function was then used, as above, to calculate differentially methylated TSSs (DMTs) in a pairwise fashion for comparison between the three sexes.

### RNA extraction and RNA sequencing for transcriptome analysis

One gonad of the pair was used for RNA extraction and sequencing; performed as described in Whiteley et al. (2021). Here repeated, RNA from isolated gonad samples was extracted in randomized batches using the Qiagen RNeasy Micro Kit (Cat. No. 74004) according to the manufacturer protocols. RNA was eluted in 14 μl of RNase free water and frozen at −80°C prior to library preparation. Sequencing libraries were prepared in randomized batches using 50 ng RNA input and the Roche NimbleGen KAPA Stranded mRNA-Seq Kit (Cat. No. KK8420). All sample RNA and library DNA was quantified using a Qubit Instrument (ThermoFisher Scientific, Scoresby, Australia), with fragment size and quality assessment using a Bioanalyzer (Agilent Technologies, Mulgrave, Australia). The libraries were sequenced on an Illumina HiSeq 2500 system at the Kinghorn Centre for Clinical Genomics (Garvan Institute of Medical Research, Sydney) in a paired-end mode with a read length of 150 bp and 25 million read-pairs per sample were obtained on average.

### Data analysis for transcriptome profiles

The transcriptomes of six samples (ZWf and ZZm samples) have been analysed and published before (Whiteley et al. 2021; Whiteley et al. 2022; Wagner et al. 2023) and have been re-analysed for this study together with the remaining ZZf samples. The paired-end RNA-seq libraries for *P. vitticeps* were trimmed using trimmomatic v0.38 (Bolger et al. 2014) with the following arguments HEADCROP:15 SLIDINGWINDOW:5:15 MINLEN:36 ILLUMINACLIPTruSeq3-PE.fa:2:30:10. Trimmed reads were then aligned to the P. vitticeps genome assembly pvi1.1 (Georges et al. 2015) using the subread-align command from Subread v2.0.2 (Liao et al. 2013) using the arguments: –sortReadsByCoordinates -t 0. Read counts were summarised to features using the Subread command featureCounts using the following parameters; -O -s 2 -p -t exon -g gene, with all others left as default. The R package edgeR v3.38.4 was used to perform differential gene expression analysis in a pairwise fashion for the three sexes as directed by the edgeR manual. Benjamini-Hochberg correction was used to correct for multiple testing with the decideTests() function with an adjusted p-value < 0.05 being qualified as a differentially expressed gene (DEG). The default edgeR normalisation method was used for differential expression analysis. For gene expression comparing genes across the sexes directly by plotting (**Figure 3B, D, F and H**) the cpm() function was used to generate counts per million (CPM). To compare expression levels of different genes within sexes the function rpkm() was used to generate reads per kilobase per million (RPKM) values for each sample. The mean RPKM value of each sex was then used to assign genes to three expression clusters with the split_quantile() function from the package fabricatR v1.0.2. Genes with no entries in RPKM dataframes were considered to have no expression or been filtered out and so, were assigned to the no expression cluster.

### Gene Ontology analysis

Gene Ontology analysis was performed using the g:GOSt function on the g:Profiler web service (Kolberg et al. 2023). As *P. vitticeps* is not a searchable organism on g:Profiler, gene lists were searched against human data to ascertain possible enrichments.

## DECLARATIONS

### Ethics approval and consent to participate

All procedures were conducted with approval from the University of Canberra Animal Ethics Committee (AEC 17-17). We confirm that all the study was reported in accordance with ARRIVE guidelines (https://arriveguidelines.org/). All the methods were in accordance with animal ethics protocols of the University of Canberra.

### Availability of data

Raw RNA and DNA bisulfite sequencing reads have been deposited in the Sequence Read Archive (SRA) repository of NCBI in BioProject PRJNA699086 (https://www.ncbi.nlm.nih.gov/bioproject/?term=PRJNA699086). Accession numbers for individual samples can be found in **Table S3.**

### Author contributions

AG and PDW led and designed the experiment. BJH and NCL performed the whole genome bisulphite sequencing. SLW carried out all other experimental procedures. SW and BJH performed the data analysis. BJH prepared the figures. BJH led the preparation of the manuscript, with SW, SLW, AG and PDW. All authors provided feedback on the manuscript and approved the final version.

## Acknowledgements

The animals used in this study were maintained and bred in the capable hands of Dr Wendy Ruscoe. We thank Dr Ira W. Deveson and James Blackburn at the Garvan Institute of Medical Research for sequencing expertise regarding the RNA sequencing. We are grateful to the members of the wildlife genetics laboratory of the Institute for Applied Ecology and the School of Biotechnology and Biomedical Sciences for the many lively discussions on elements presented in this paper.

## SUPPLEMENTARY FIGURE LEGENDS

**Figure S1. CpG read depth distribution for DNA bisulphite sequencing.**

Histograms of the cumulated read depth for all CpGs in a 1kb region. The percentage of regions with a cumulated CpG read depth of at least 20 is given and marked by the black vertical line. Only those regions were included in the further analysis. The histograms are shown for all replicates. ZWf: concordant female, ZZf: sex reversed female, ZZm: concordant male.

**Figure S2. CpG methylation rate.**

Histograms of the methylation rate of all CpGs in a 1kb region. Only regions with a cumulated CpG read depth of at least 20 are included. The histograms are shown for all replicates. ZWf: concordant female, ZZf: sex reversed female, ZZm: concordant male.

**Figure S3. Methylomes of the three sexes**

Analysis of methylomes of the three sexes in 1kb regions with a CpG read depth of at least 20 in all eight samples. (A) A dendrogram for the methylation rate of 1 kb regions. Methods used for distance and clustering are ‘correlation’ and ‘ward’, respectively. (B) a PCA plot of the methylation rate of 1 kb regions. (C)-(E) Volcano plots of differential methylation of 1 kb regions with for each iteration of a pairwise comparison of all sexes. Points are coloured according to their corrected p (q) value with the following groupings: q>0.01, q<0.01 and q<0.001.

**Figure S4. Transcriptomes of the three sexes are distinct.**

A principal component analysis (PCA) of the RNA sequencing read count per gene. The replicates of each sex, ZZf, ZWf and ZZm cluster together but separately from other sexes.

## SUPPLEMENTARY TABLES

**Table S1. Gene Ontology of differentially expressed genes with differentially methylated TSSs, ZWf vs ZZf.**

**Table S2. List of differentially expressed genes with differentially methylated TSSs.**

DEGs with DMTs are shown for all comparisons: ZWf-ZZm, ZZf-ZWf, and ZZf-ZZm.

**Table S3. Sample information and accession.**

