## Supplementary figures and images for "DNA methylation regulates a key sex differentiation gene in *Pogona vitticeps*, a dragon lizard with sex reversal"

### Figure S1

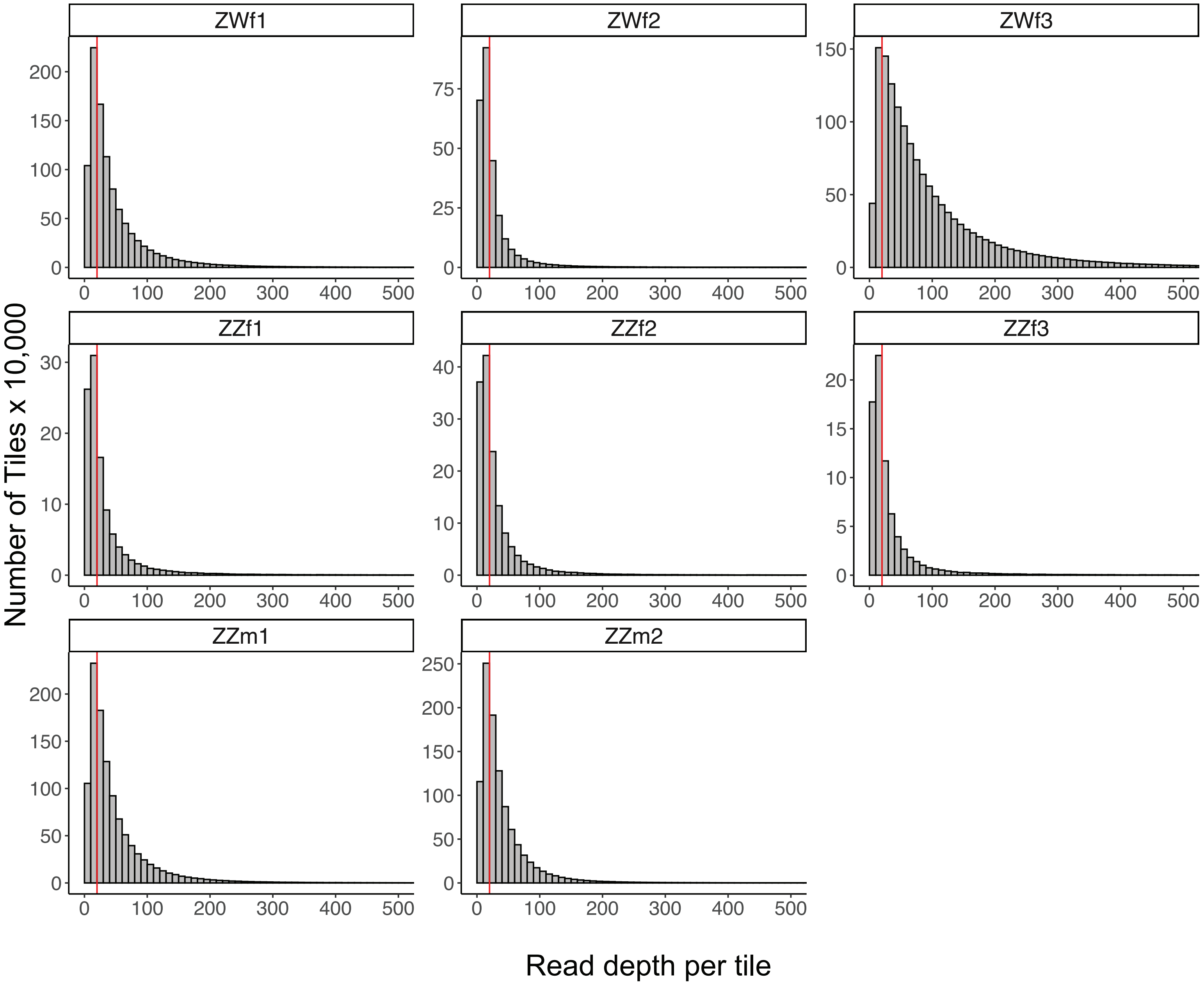

### Figure S2

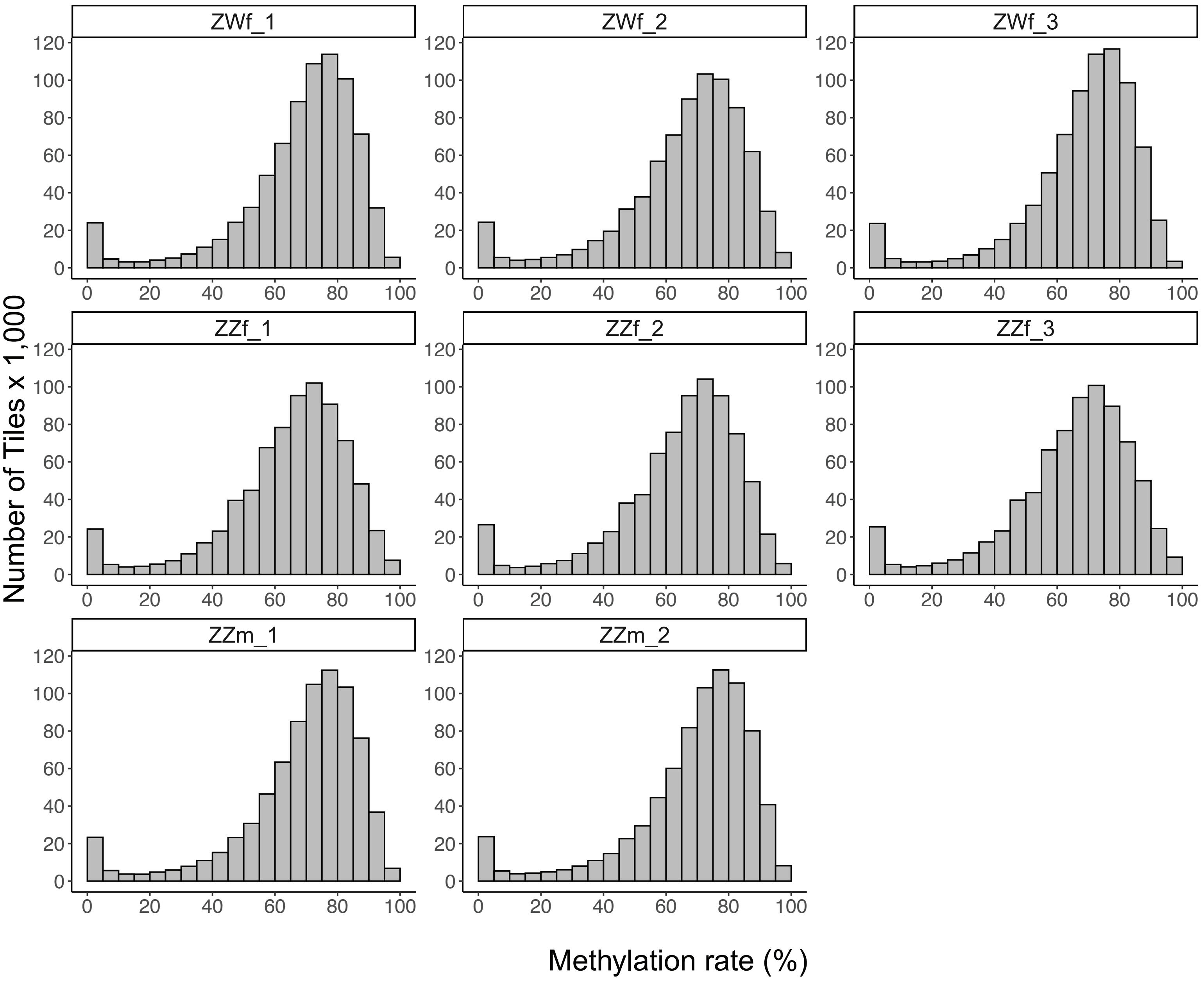

### Figure S3

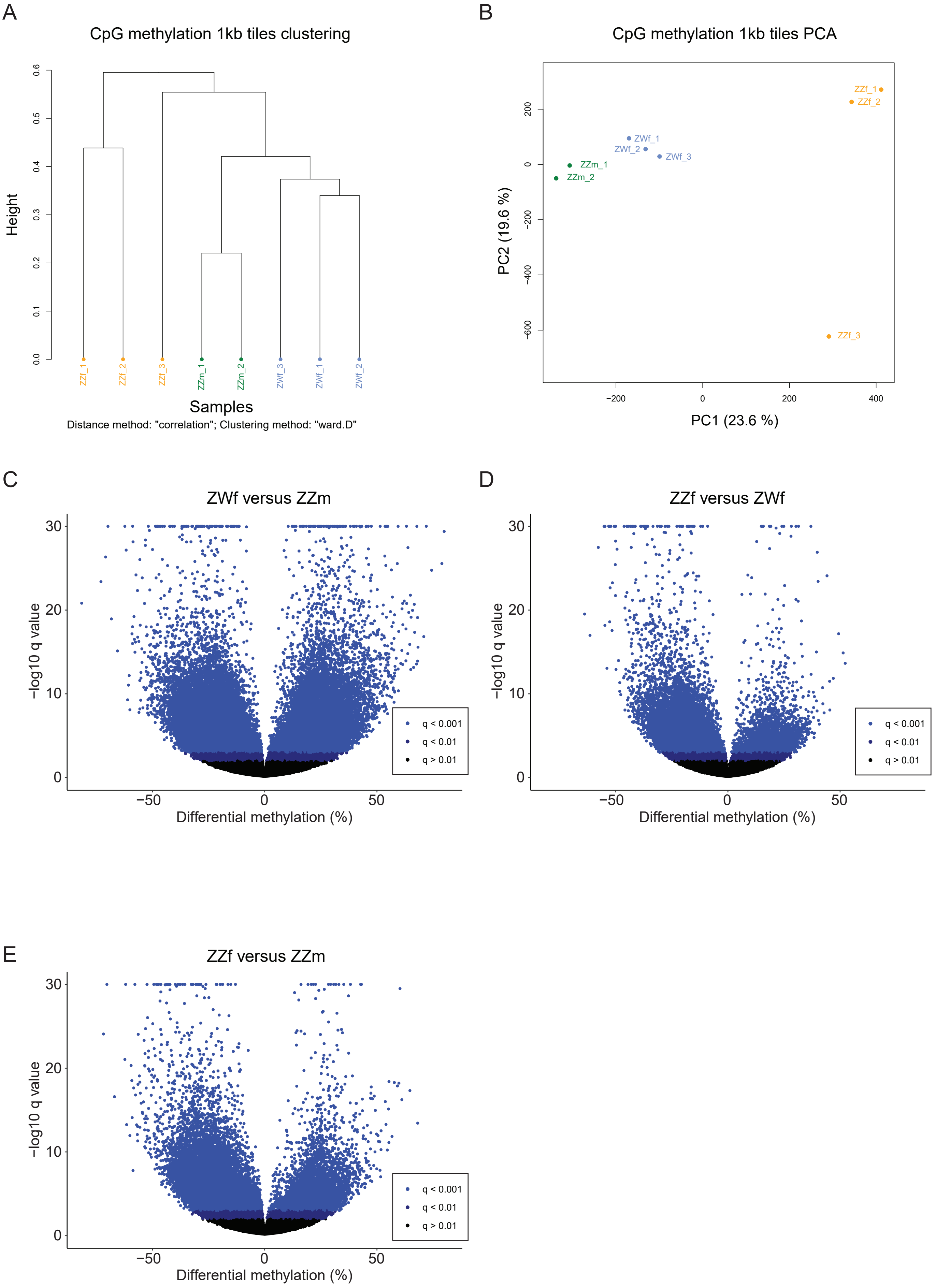

### Figure S4

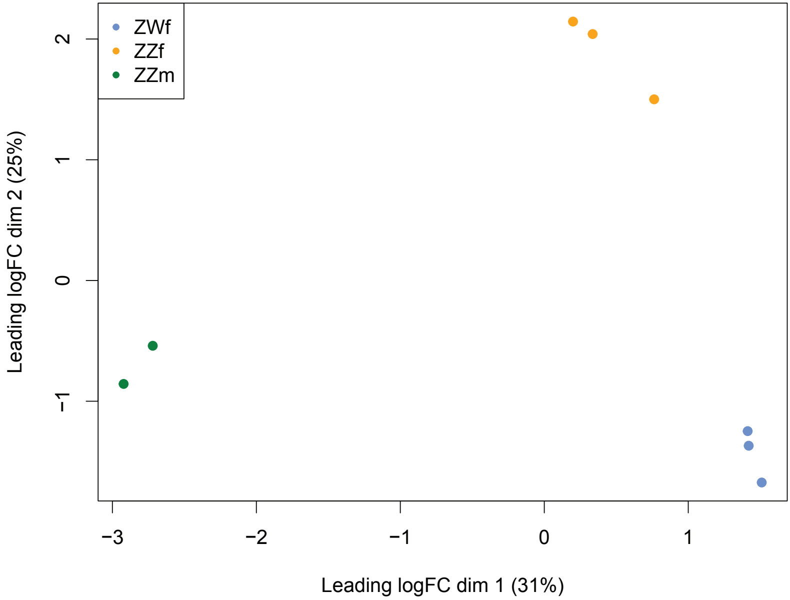
